# CASP15 cryoEM protein and RNA targets: refinement and analysis using experimental maps

**DOI:** 10.1101/2023.08.07.552287

**Authors:** Thomas Mulvaney, Rachael C. Kretsch, Luc Elliott, Joe Beton, Andriy Kryshtafovych, Daniel J. Rigden, Rhiju Das, Maya Topf

## Abstract

CASP assessments primarily rely on comparing predicted coordinates with experimental reference structures. However, errors in the reference structures can potentially reduce the accuracy of the assessment. This issue is particularly prominent in cryoEM-determined structures, and therefore, in the assessment of CASP15 cryoEM targets, we directly utilized density maps to evaluate the predictions. A method for ranking the quality of protein chain predictions based on rigid fitting to experimental density was found to correlate well with the CASP assessment scores. Overall, the evaluation against the density map indicated that the models are of high accuracy although local assessment of predicted side chains in a 1.52 Å resolution map showed that side-chains are sometimes poorly positioned. The top 136 predictions associated with 9 protein target reference structures were selected for refinement, in addition to the top 40 predictions for 11 RNA targets. To this end, we have developed an automated hierarchical refinement pipeline in cryoEM maps. For both proteins and RNA, the refinement of CASP15 predictions resulted in structures that are close to the reference target structure, including some regions with better fit to the density. This refinement was successful despite large conformational changes and secondary structure element movements often being required, suggesting that predictions from CASP-assessed methods could serve as a good starting point for building atomic models in cryoEM maps for both proteins and RNA. Loop modeling continued to pose a challenge for predictors with even short loops failing to be accurately modeled or refined at times. The lack of consensus amongst models suggests that modeling holds the potential for identifying more flexible regions within the structure.

## 1. Introduction

Assessment of models in CASP is traditionally based on comparing predicted coordinates with the coordinates of reference structures provided by experimentalists. For evaluation purposes, the experimental structures are considered the ‘gold standard’. However, experimental structures by their nature are only models themselves – their construction involves a certain degree of subjectivity in interpreting density maps and translating them to atomic coordinates. In several previous CASPs, in parallel to the coordinate-to-coordinate evaluation, we carried out an evaluation of models versus the experimental data for a subset of cryoEM-derived structures ^1, 2^, where experimental uncertainty was expected to be larger than that in X-ray structures. In this article, we continue this trend and check the fit of some of the best CASP15 models to cryoEM density maps. We also study how the density-guided refinement of these models improves their fit to map, and how the refined models fare with regards to the experimental structures. For the first time, besides the protein targets, we analyze RNA structures.

The number of structures newly solved by 3D-EM roughly doubles every two years and totals 14,500 as of March 2023, constituting more than 8% of protein structures in the whole PDB (http://www.rcsb.org/) ^3^ (compared to around 4% only two years ago). Reflecting this growth, CASP also registered an uptick in the percentage of cryoEM targets. In CASP14, 7 out of 54 evaluated targets (13%) were determined by cryoEM, while in CASP15 the corresponding numbers were 27 out of 93 (29%), including 8 of the 12 (67%) RNA-containing cryoEM structures.

While AlphaFold2 did not participate in the assembly category in CASP14, it was noted that its predictions could have alleviated many interface modeling errors ^4^. Since then, AlphaFold-Multimer, RosettaFold ^5^ and AF2Complex ^6^ are a few examples of a growing number of deep-learning approaches to complex prediction. Predictions of oligomeric targets were sufficiently good in CASP15 to directly refine whole proteins and complexes rather than smaller evaluation units (as done in previous studies). To test the applicability of the predictions in real-world cryoEM structure determination tasks, we not only examined the refinement of complete complexes but also investigated a method for identifying good candidates for refinement. Additionally, given the improvement in the average cryoEM map resolution, we decided to not only refine the best-predicted models into the corresponding maps but also assess higher resolution aspects of predicted models, such as their side-chain orientations.

Whilst cryoEM has been an important method for studying proteins, often at near-atomic resolution (see H1114), cryoEM experiments have not yet been able to achieve the same levels of resolution for RNA-only structures. The lower resolution nature of the data makes RNA structures ideal test cases for flexible-fitting into cryoEM maps based on *de novo* models, where possible motions of the structure can be calculated from an initial structural model and the experimental data. However, structure prediction for RNA is far less mature than for proteins, making RNA refinement into cryoEM maps particularly challenging.

## 2. Materials and Methods

### 2.1 Selection of models for refinement from proteins and protein complexes targets

In CASP15, with the accuracy of domain modeling expected to be on a par with or better than in previous years, we focussed on the fitting and refinement of multidomain and oligomeric models **(Fig. 1**, **Table 1)**. Protein models were selected by combining two approaches **(Fig. 2)**. In the first approach, we used the following CASP metrics to define accurate models: all predictions required an lDDT (lDDTo for oligomers) score greater than 0.7. Additionally, predictions for monomeric targets required a GDT_TS score greater than 0.7. In the case of oligomeric targets, predictions with QS, TM and F1 scores ^4, 7, 8^ all greater than 0.7, 0.8 and 0.6 respectively were eligible for refinement. Often in an experimental setting, little is known about the target structure and therefore the first step is to fit the structure in the map using a global search. We therefore, as a “control” experiment, also chose the prediction for each of the above targets **(Fig. 1**, **Table 1)** which ranked best against the experimental map **(see 2.2)**.

**Fig. 1:**
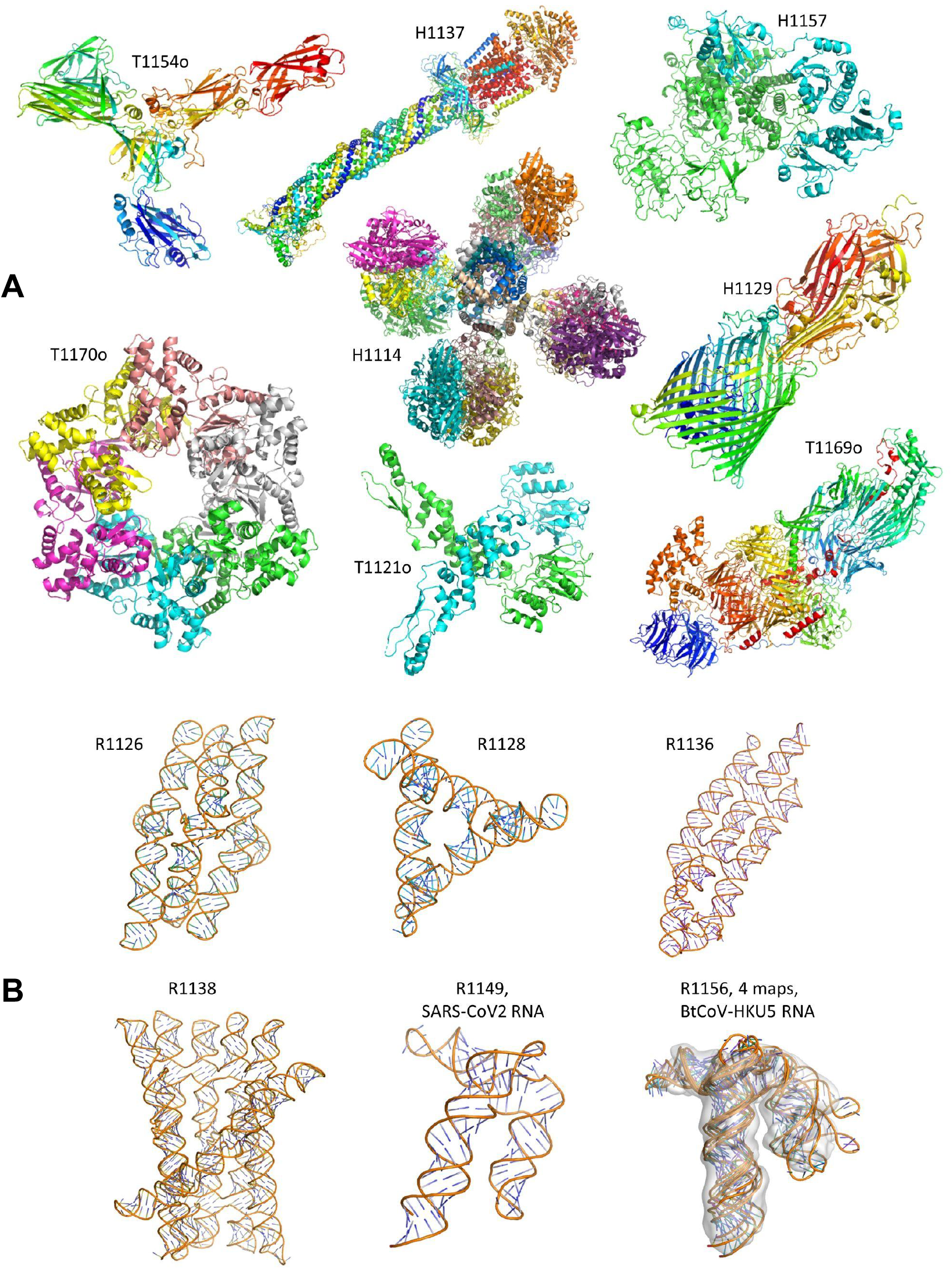
Overview of the Cryo-EM targets used for refinement and analysis in CASP15: Reference structures for 8 protein targets (A) and 6 RNA targets (B) solved by cryoEM in CASP15.

**Fig. 2:**
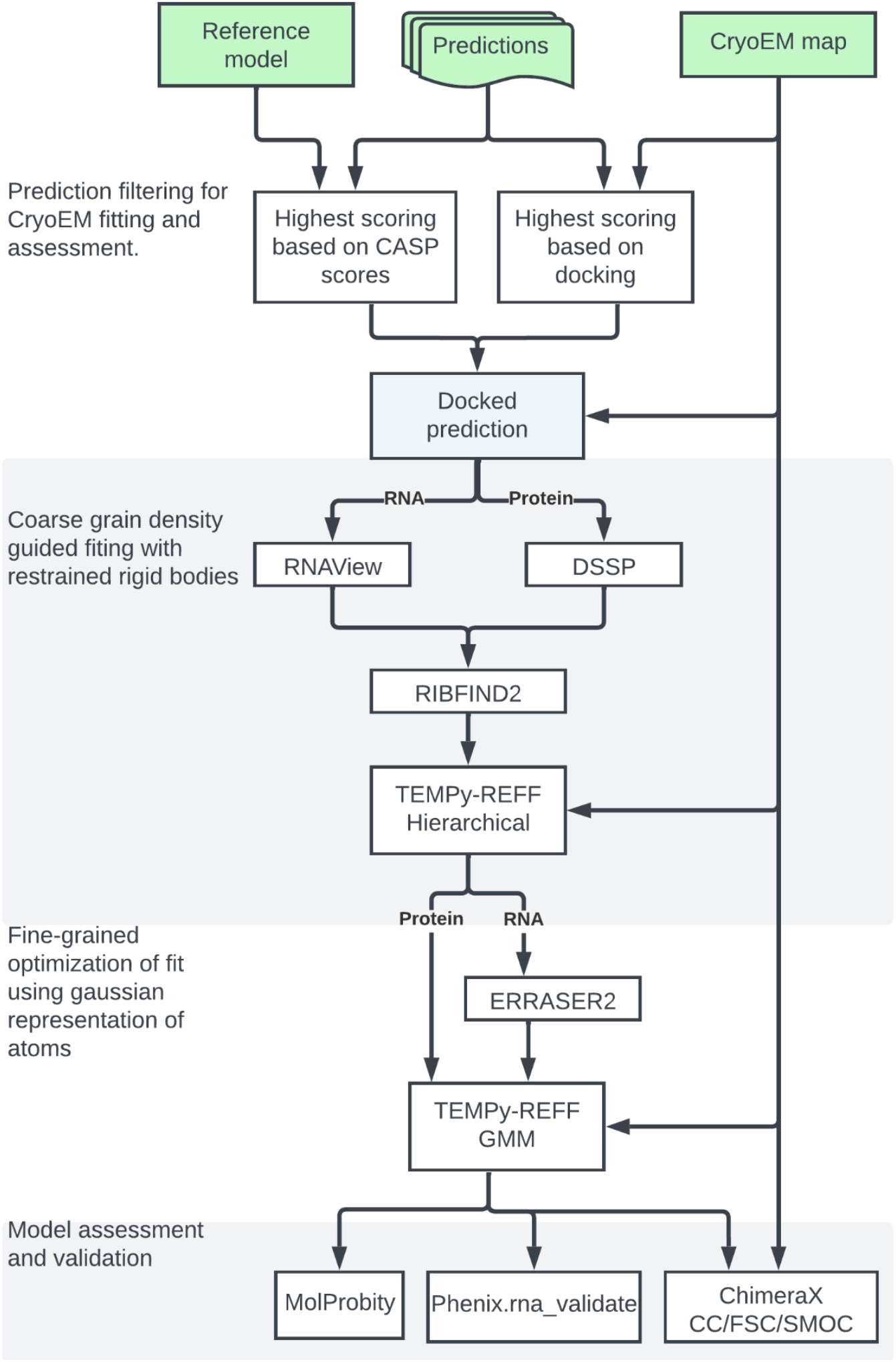
Docking and refinement pipeline. Submitted predictions were chosen for cryoEM refinement based on CASP metrics (using the reference structure) and using a ranking scheme which assesses the overall fit of predictions constituent chains to the experimental map. Selected models are then flexibly fit to the experimental data using a coarse-grained hierarchical fitting protocol. Corrections were made to RNA model geometry using ERRASER2 followed by an atomistic refinement scheme based using TEMPy-REFF. Models were then assessed using MolProbity, Phenix.rna_validate, ChimeraX CC and TEMPy SMOC scores.

**Table 1.**
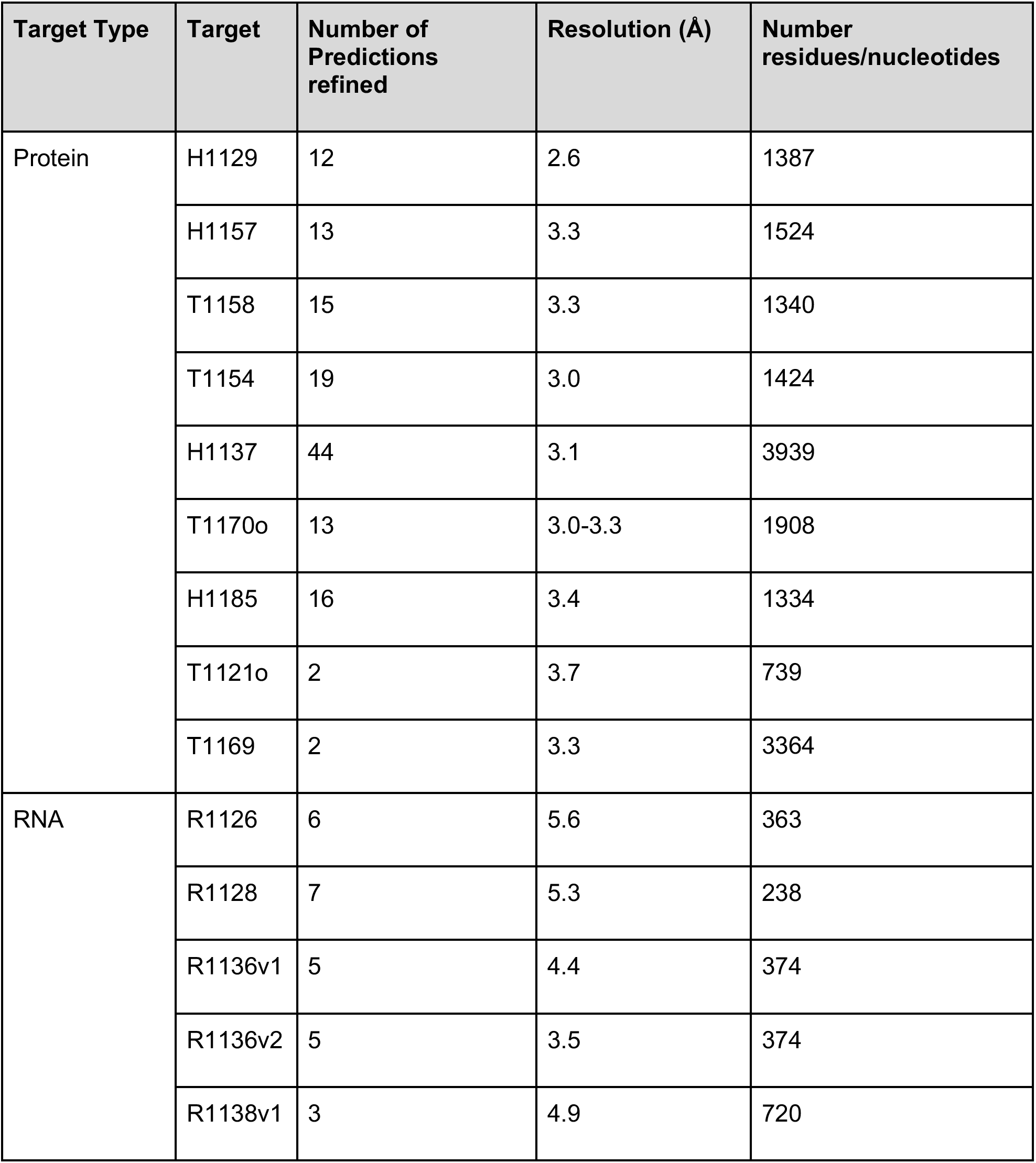
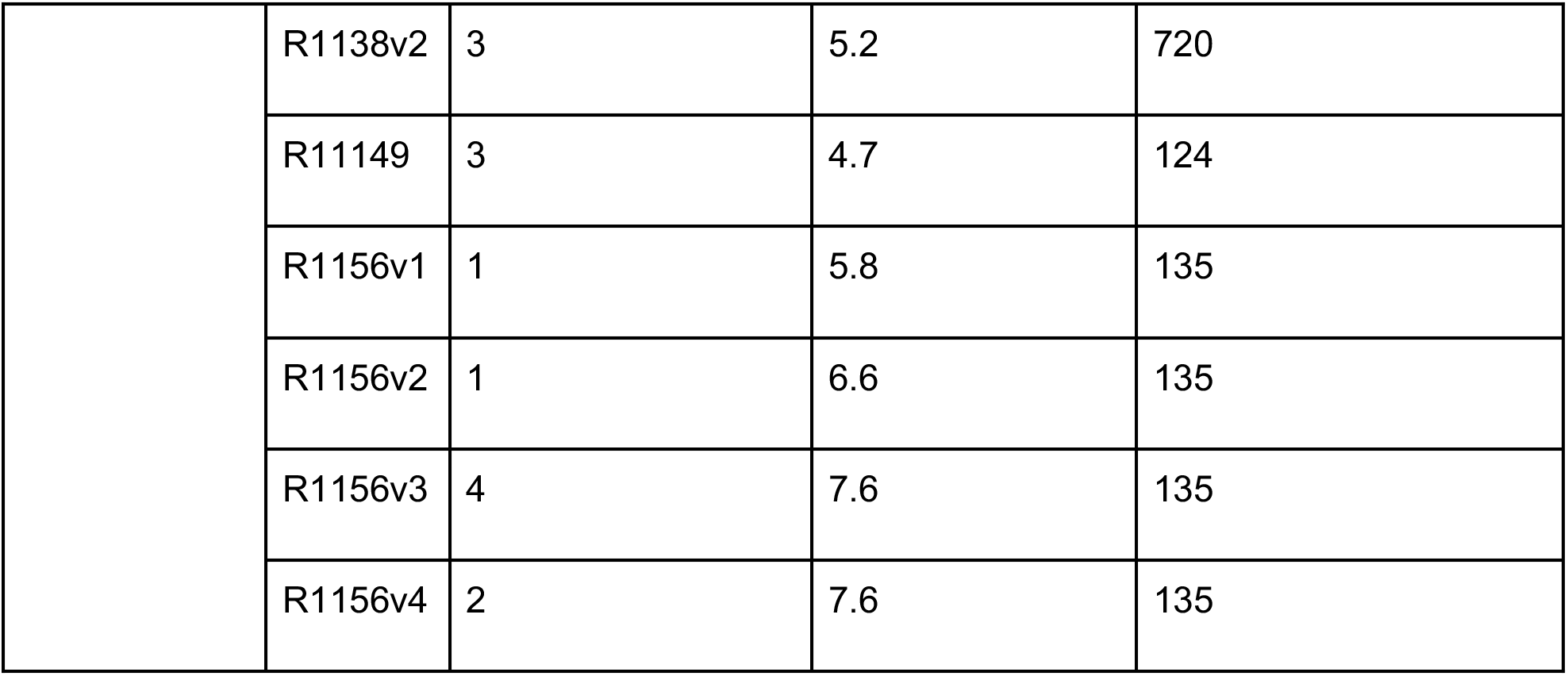
Overview of targets. Targets with predictions which met the minimum score criteria were refined.

| Target Type | Target | Number of Predictions refined | Resolution (Å) | Number residues/nucleotides |
| --- | --- | --- | --- | --- |
| Protein | H1129 | 12 | 2.6 | 1387 |
|  | H1157 | 13 | 3.3 | 1524 |
|  | T1158 | 15 | 3.3 | 1340 |
|  | T1154 | 19 | 3.0 | 1424 |
|  | H1137 | 44 | 3.1 | 3939 |
|  | T1170o | 13 | 3.0-3.3 | 1908 |
|  | H1185 | 16 | 3.4 | 1334 |
|  | T1121o | 2 | 3.7 | 739 |
|  | T1169 | 2 | 3.3 | 3364 |
| RNA | R1126 | 6 | 5.6 | 363 |
|  | R1128 | 7 | 5.3 | 238 |
|  | R1136v1 | 5 | 4.4 | 374 |
|  | R1136v2 | 5 | 3.5 | 374 |
|  | R1138v1 | 3 | 4.9 | 720 |
|  | R1138v2 | 3 | 5.2 | 720 |
|  | R11149 | 3 | 4.7 | 124 |
|  | R1156v1 | 1 | 5.8 | 135 |
|  | R1156v2 | 1 | 6.6 | 135 |
|  | R1156v3 | 4 | 7.6 | 135 |
|  | R1156v4 | 2 | 7.6 | 135 |

### 2.2 Selection of models for refinement based on ranking of individual protein chains

Instead of rigidly fitting the entire complex in the map, one can identify the optimal initial position for each of the protein components in the model using an exhaustive search or another heuristic. Predictions were re-ranked based on this global fitting approach using Cross-Correlation (CC).

The docking of models in this study was carried out using two automatic docking programs, Molrep ^9, 10^ and PowerFit ^11^. Both programs use a six-dimensional search to maximize an overlap-correlation score between a given model and the map file. Molrep incorporates a Spherically Averaged Phased Translation Function (SAPTF), followed by a Rotation Function (RF) and Phased Translation Function (PTF), which achieves a suggested first fit and then improves the overlap score with a six-dimensional optimisation search ^9, 10^. On the other hand, PowerFit incorporates an exhaustive six-dimensional search, including rotation at a pre-set angle sampling density and translation across the map file. Input parameters for the docking included the input map file, model and resolution ^11^. The top model was determined by the CC score calculated using ChimeraX ^12^.

A group ranking was generated as follows using the complete *chain* submissions submitted by groups instead of the CASP-defined Evaluation Units (EUs) ^13^. Predictors may submit five models for each target. To reduce the computational time required for the docking process, only the first submitted model for each target per group was considered. For each target, a score was assigned per group reflecting its position in the CC ranking for that target. The top model was given a score of 123 since this was the total number of groups. An automatic rank of 0 was given where a group did not submit a prediction for a given target. For an overall group ranking, a cumulative score for each group was tallied across all targets for which that group submitted a prediction. For comparison, similar rankings were done for each group and target using the composite S_CASP15_ score defined by ^14^

For single chain targets, the prediction from the top group was chosen as the starting candidate. For oligomeric targets (H1114, H1129, H1158, T1121o, T1170o, H1185), a cumulative score of the individual chains was tallied. The model from the highest scoring group across all chains for a target was selected for refinement. For these models, no attempt was made to recombine individual fitted chains: instead the originally submitted multi-chain assembly was re-docked so that this full assembly was the starting model for the refinement process.

### 2.3 Selection of models for refinement from RNA targets

All RNA-containing cryoEM targets were considered for refinement. If there were multiple experimental maps, predicted models were selected separately for each map. The predictors were not asked to predict these conformations separately and hence, in some cases, the same predicted model was refined against multiple maps. Due to the prediction accuracy, there were a limited number of models that fit well in each map, so all models submitted by each team were considered. The best models were selected as the top ranked structures across all submitted models based on the previously described map-to-model Z-score, Z_EM_ ^15^. Due to the fit qualities an automatic threshold would result in few models per target, so manual visual inspection was additionally used, to select models that, even without good fits, we thought were the most promising for refinement. Based on these rankings and visual inspection of fit of the top 10 ranking models by an expert, a final set of models for each target were selected.

### 2.4 Model fitting and refinement

As per the CASP13 and CASP14 modeling experiments ^1, 2^, predictions of cryoEM targets were positioned in the density by aligning them to the target and then optimizing local fit to density before refinement using ChimeraX fit-to-map function. The fitting and refinement stages of the pipeline **(Fig. 2)** used a hierarchical approach built on our previous protocol designed for Flex-EM/RIBFIND ^16^. This approach was previously shown to allow large conformational changes to take place and avoid trapping parts of the model in small density pockets during fitting. The TEMPy-REFF software package ^17^ supports this hierarchical approach by automatically breaking down clusters as refinement progresses. This package offers a number of force-fields and routines for building pipelines for fitting and refinement using the OpenMM molecular dynamics engine ^18^. In our application of the software here **(Fig. 2),** the models were iteratively broken down into smaller rigid-bodies using the RIBFIND2 software [https://ribfind.topf-group.com/]. These subunits were composed of interacting secondary structure elements determined using DSSP ^19^ for proteins and RNAView ^20^ for RNA. Each subunit was then subjected to a fitting force which is simply the negative gradient of the cryoEM map ^21^, whilst strong harmonic restraints maintain the overall geometry of the subunit.

In the last stage, the models were refined using a Gaussian mixture model (GMM)-based potential similar to the component-based approach of ^22^, but this time applied to atoms. In this scheme, each atom is represented by a Gaussian where the mean is given by the atom’s position, sigma can intuitively be thought of as its resolution or Bfactor, and the height is the atomic number which approximates the scattering potential of neutral atoms. Using the expectation-maximization algorithm for GMMs, the atomic positions are successively updated so as to maximize the likelihood that the Gaussians represent the experimental data while at the same time an AMBER14 force-field ^23^ corrects stereo-chemistry.

Despite the RIBFIND rigid body restraints, early testing of the pipeline showed that some RNA models suffered from distortions. Therefore, our final protocol included an additional stage of corrections using the ERRASER2 program, an unpublished accelerated version of the ERRASER protocol for the refinement of RNA in crystallographic and cryoEM maps ^24^. It is available as part of the ROSETTA package ^25^. However, here ERRASER2 was used only for optimisation of the structure and not the fit to the density before final refinement of the structure using the GMM (see Listing 1).

### 2.5 Model assessment measures for protein models

The protein predictions for cryo-EM protein targets and the subsequent refined models were evaluated for their goodness-of-fit to the experimental cryo-EM density map (model-to-map goodness-of-fit) using the following metrics: The local (per-residue) goodness-of-fit was evaluated with the TEMPy2 Segmented Manders’ Overlap Coefficient (SMOC) score ^16^ and global goodness-of-fit using the ChimeraX cross-correlation measurement. The SMOC score represents the Manders’ overlap coefficient for overlapping residue fragments: it is computed on local spherical regions around the seven residues in the current window. Overlapping windows are used, producing one numerical value per residue. SMOC scores can be calculated for the whole structure by averaging the per-residue scores. In order to compare the quality of fit to the density of side-chain *vs.* backbone, we have implemented two new “localised” SMOC scores in TEMPy: SMOCs and SMOCb. These scores assess the voxels around the side-chain atoms (SMOCs) and around the backbone atoms (SMOCb), respectively. To compute the SMOCs and SMOCb scores, each residue from the predictions was locally aligned to the target using the C-alpha atoms of the residue and its immediate neighbors. Because side-chains are a high-resolution feature, we did not use sliding windows in this case, i.e., SMOCs and SMOCb scores were computed on the aligned residues. The geometry of the targets, the predictions and the refined models were all assessed using MolProbity ^26^.

### 2.6 Model assessment measures for RNA models

For CASP15, we have implemented a new SMOC score in TEMPy – SMOCn – to assess the fit of nucleic acid chains. SMOCn is calculated similarly to the original SMOC score, which was designed to assess the protein chain in the density, by sliding windows around nucleotides instead of amino acids ^16^. Due to resolution limitation, the “localized” SMOCb and SMOCs scores were not used for RNA. As the RNA experimental maps were generally of a lower resolution than their protein counterparts, assessing geometry was important to ensure models were not overfit to the maps. RNA Validate, which is part of the Phenix ^27^ software package, was used to assess the geometry of the RNA targets, predictions and refinements. We focussed our geometry analysis on the ‘average suiteness’ scores produced by RNA Validate. ‘Suites’ are defined by the pucker of two consecutive backbone sugars and the five torsion angles between them. Empirical studies have shown that these suites inhabit a number of characterized states in 7-dimensional space. ‘Average suiteness’ is a measure of how well the suites in an RNA model match the discrete conformers found in the empirical data ^28^.

## 3 Results

### 3.1 Protein targets: ranking and selection

#### 3.1.1 Selection of protein targets for refinement using CASP criteria

We refined 136 predictions for multi-domain proteins and protein complexes **(Table 1)** that either passed our filter based on CASP score (see 2.1) or ranked first based on the fit of individual chains (see 2.2). For most targets, there was an overlap between the two, i.e., the top-ranked model based on global fitting of the chain/s was included in the list of models which had good CASP scores.

The only target listed which did not have models that passed the CASP based selection criteria was T1169. Predictions of individual domains in T1169 were good but the full protein models were not accurate enough to pass the threshold due to partially inaccurate domain organization. This protein was the largest single chain model in CASP history with 5 domains and over 3000 residues. Here we chose the model with the highest GDT_TS score (GDT_TS=57.7, lDDT=0.63) which was from Yang-server (group 229). Finally, we did not refine predictions for target H1114 for which the corresponding cryoEM map is at 1.52 Å resolution. Given the high resolution of the map and the high quality of the predictions for this target (the best model had a TM-score=0.97, olDDT=0.86, QS-score=0.79, F1-score=84.13), we decided to use it for side-chain analysis instead.

#### 3.1.2 : Choice of docking software for chain ranking

A Spearman rank correlation coefficient was performed to compare the rankings produced by Molrep and PowerFit. For the majority of cryoEM targets there were no significant differences between the PowerFit and Molrep rankings. However, for a minority of cryoEM targets, significant differences were seen. The greatest differences were for T1114s3 (PowerFit median CC: 0.6483, Molrep median CC: 0.587), T1137s1 (PowerFit CC: 0.17285, Molrep CC: 0.12070), T1137s3 (PowerFit CC: 0.1894, Molrep CC: 0.1273), T1137s5 (PowerFit CC: 0.20345, Molrep CC: 0.11065), T1157s1 (PowerFit CC: 0.46999’, Molrep CC: 0.6101) and T1185s1 (PowerFit CC: 0.317, Molrep CC: 0.439). Since PowerFit outperformed Molrep in those cases more often than the reverse, PowerFit’s placements were chosen for the rankings.

#### 3.1.3 Chain rankings

There was a significant, strong positive correlation between the cumulative S_CASP15_ rankings and the cryoEM-based docking rankings **(Fig. 3)**. The top five groups from the docking rankings, in order, were: Yang, BAKER, GuijunLab-Assembly, FoldEver and PEZYFoldings. Each of these groups submitted predictions for each target with the Yang group ranking consistently high across all targets. Additionally, Yang had the most (three) top ranking models by the docking rankings **(Table 2)**. Each of the top groups used different flavors of AlphaFold 2 for the predictions with the exception of BAKER who used RosettaFold. For making comparisons in performance, control representations of AlphaFold 2 are annotated **(Fig. 3)** with group names NBIS-af2-multimer, NBIS-af2-standard, Colabfold and Colabfold_human. Colabfold and Colabfold_human submitted predictions for every target but their results, while confirming the value of these readily available predictions for cryo-EM map fitting, were not among the very best. The best ranked prediction for each target was selected for refinement if it was not already selected based on CASP criteria **(see 3.1.1).** These are listed in **Table 3**. H1137 was excluded from the ranking as the CC scores for each chain failed to produce consistent results.

**Fig. 3.**
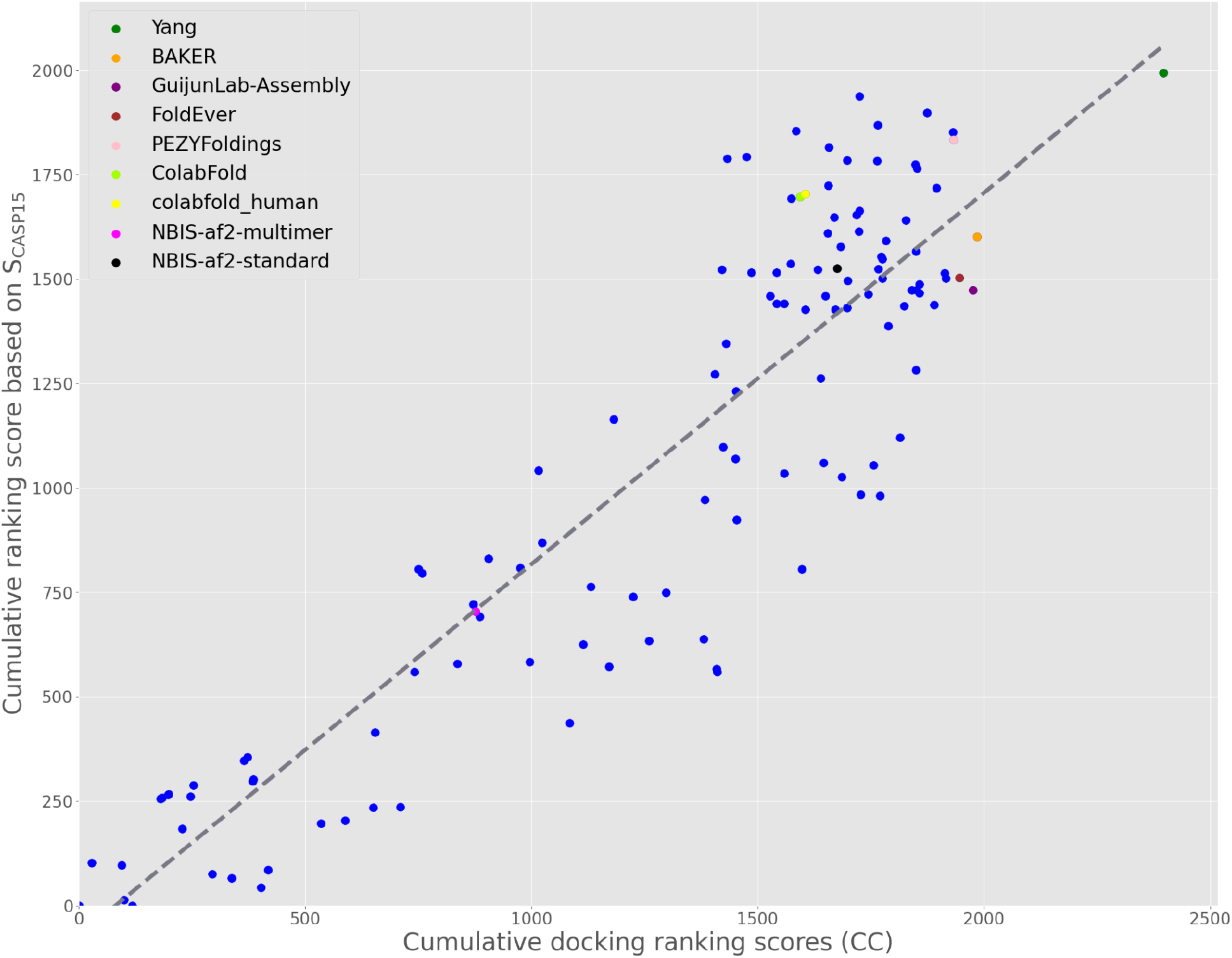
Group ranking for cryoEM targets. Cumulative per-group docking ranking scores plotted against S_CASP15_ rankings across docking targets where S_CASP15_ scores were available (oligomeric reference structures were split into individual chains – see also Table 2). The gray line indicates the line of best fit with a strong positive correlation between the two rankings (r=0.827, p<0.0001). The top five performing docking ranking groups are labeled, as are the ‘control’ AlphaFold 2 submissions.

**Table 2:**
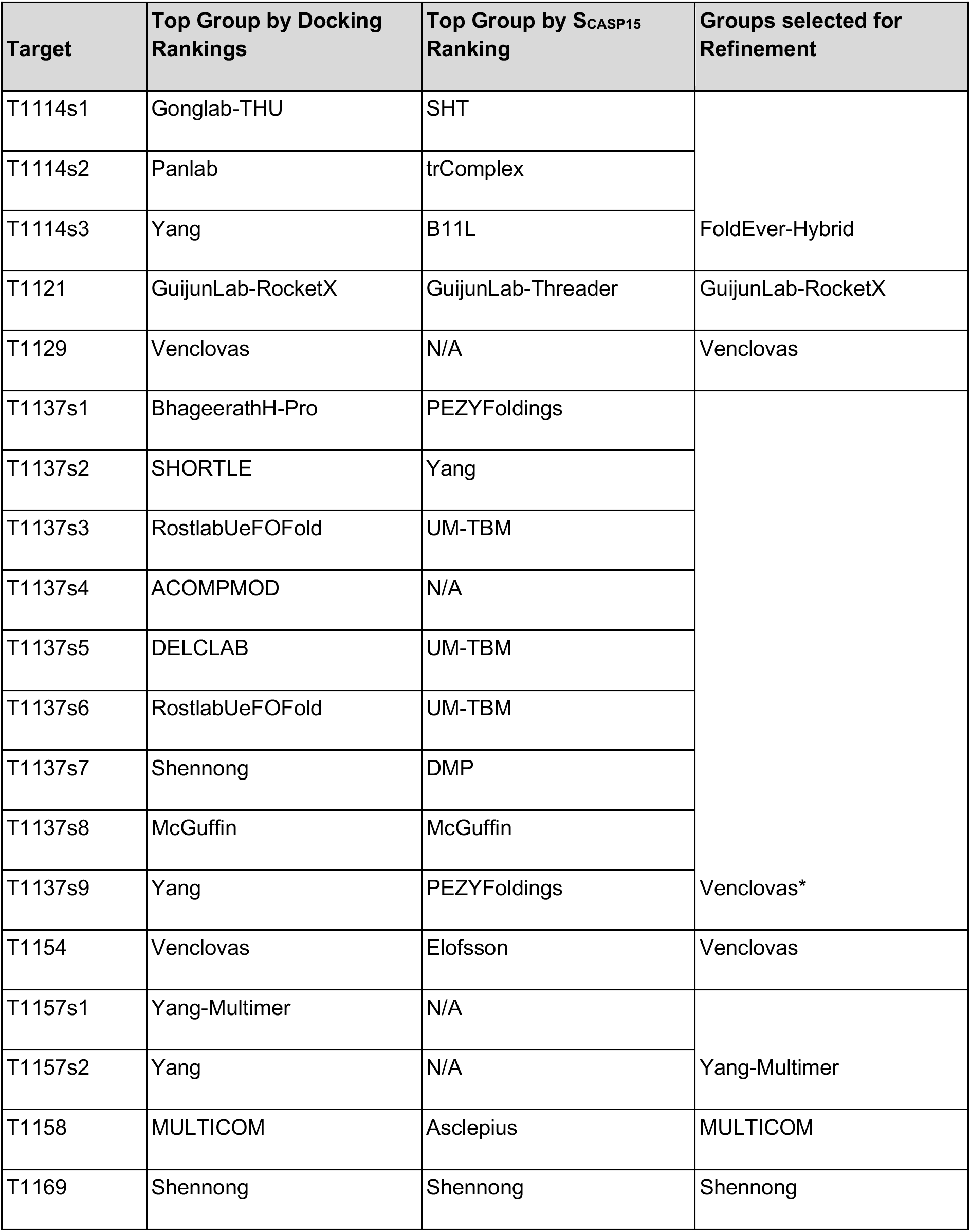
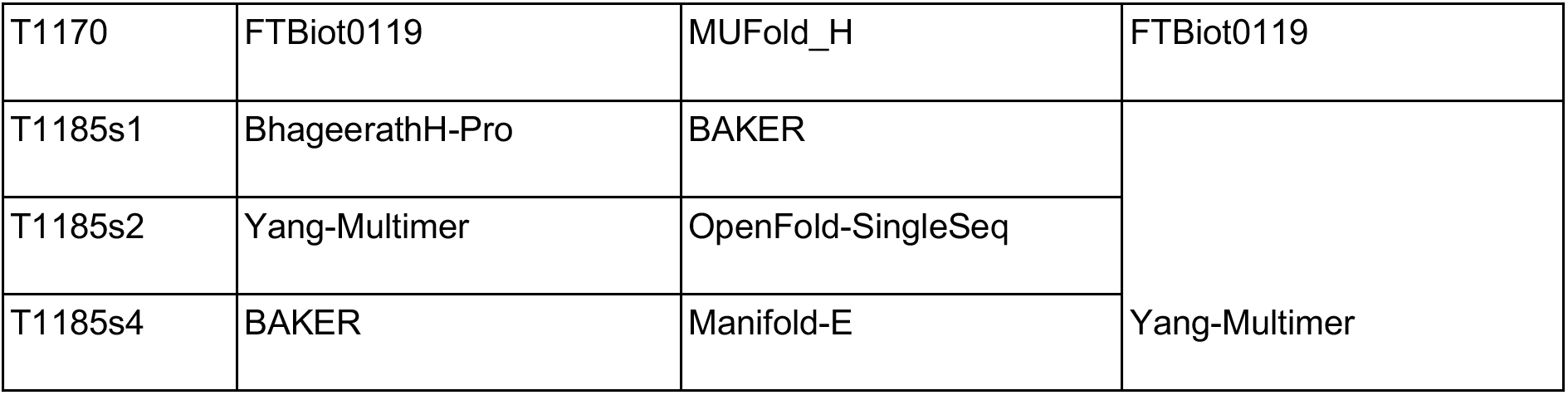
Group ranking based on docking. The top-scoring CC model for each target. Also indicated are the top-scoring groups for the same targets, in the general CASP assessment using the CASP15 score ^14^. Some chain models did not receive a CASP15 score because certain elements used in the CASP15 score formula were not calculated since the chain in question was split into multiple AUs. These were given an N/A classification. *These targets were selected in a different way – see section 3.4.3

**Table 3:** Predictions found only by the docking-based ranking method: Values for TM, GDT-TS, (o)lDDT and QS scores of predictions and their respective refined models. TM and QS scores were not applicable to monomeric targets.

| Target | Group | (o)IDDT |  | GDT-TS score |  | QS-score |  | TM score |  |
| --- | --- | --- | --- | --- | --- | --- | --- | --- | --- |
|  |  | Pred | Ref. | Pred | Ref. | Pred. | Ref. | Pred. | Ref. |
| T1154 | 494 | 0.84 | 0.77 | 0.63 | 0.78 | NA | NA | NA | NA |
| T1158 | 367 | 0.84 | 0.86 | 0.64 | 0.92 | NA | NA | NA | NA |
| T1169 | 466 | 0.66 | 0.67 | 0.53 | 0.72 | NA | NA | NA | NA |
| H1157 | 239 | 0.66 | 0.7 | NA | NA | 0.67 | 0.8 | 0.78 | 0.80 |
| T1170o | 165 | 0.70 | 0.85 | NA | NA | 0.61 | 0.85 | 0.90 | 0.90 |
| T1121o | 091 | 0.74 | 0.69 | NA | NA | 0.31 | 0.25 | 0.66 | 0.64 |

### 3.2 Protein targets – refinement of top predictions

#### 3.2.1 Overall model analysis

Average SMOC scores of predictions prior to refinement were poor with a large degree of variation among the predictions for each target **(Fig. 4A)**. After refinement, average SMOC scores were closer to those of the respective targets, typically with significantly reduced variance. For example, the top models refined from the predictions of target T1154 had a SMOC curve very similar to that of the target SMOC curve **(Fig. 4B, C)**. Interestingly, all top predictions for this target based on CASP criteria could be refined in the N-terminal part of the structure, despite its initial wrong orientation. This is likely to be attributed to the hierarchical refinement protocol, where the N-terminal is first pulled into the density as one rigid body. On the other hand, in the regions of residues 810-814 **(Fig. 5B)**, there is a sharp drop in the SMOC plot due to the “loopy” characteristics of the region (see below). In fact, most targets had some loops which did not reach the high SMOC scores seen in the rest of the structure after refinement, suggesting these regions were poorly modeled and bringing down the average SMOC scores. Specific cases are explored in detail in **(see 3.2.2)**.

**Fig. 4:**
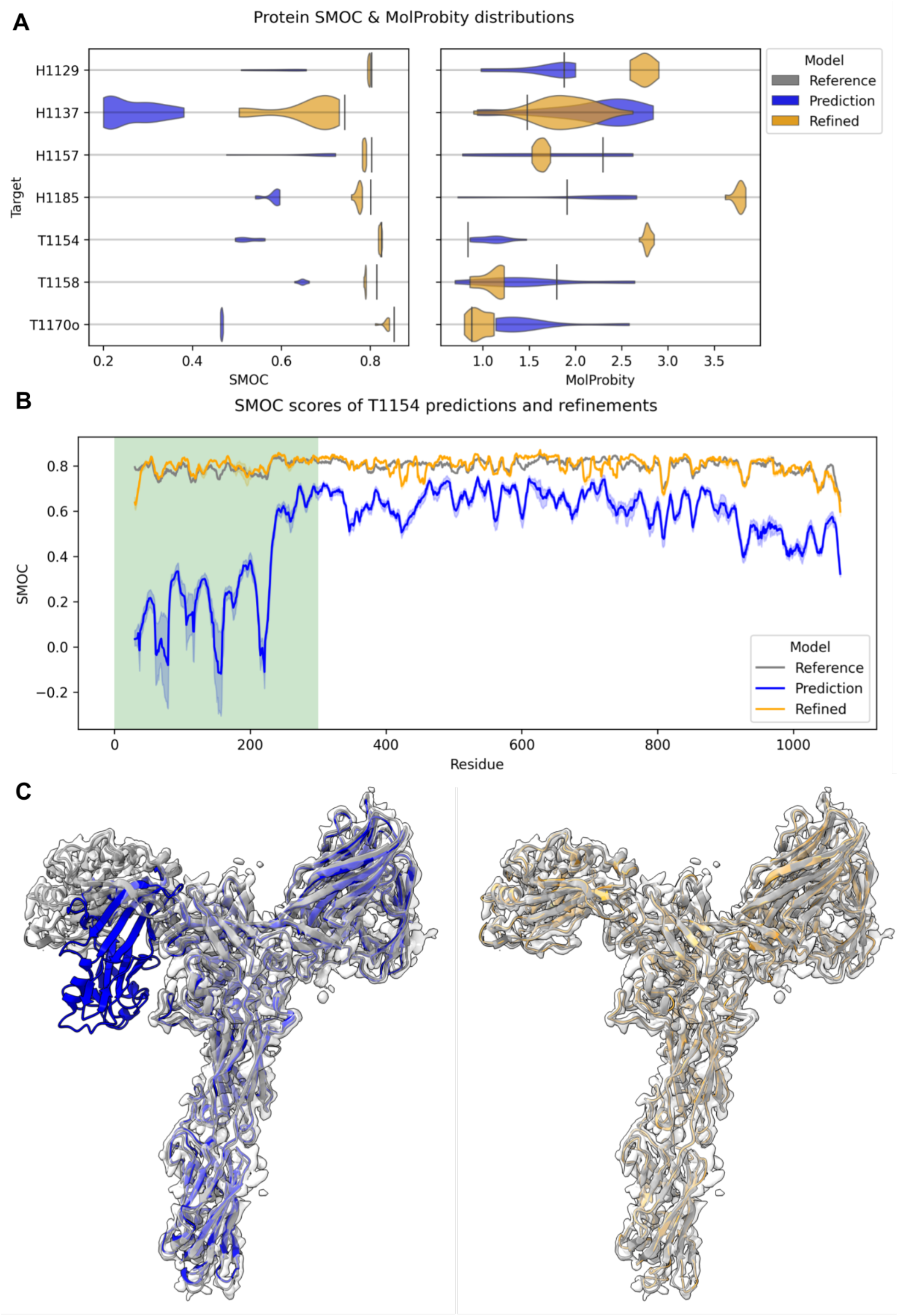
Overview of protein refinement results. In **(A)**, the distribution of average SMOC scores for each prediction is shown before and after refinement with respect to the target model. In **(B)**, the residue level SMOC plot is shown for T1154 and its predictions. The dark orange and blue lines are the mean refined and docked SMOC scores with the minimum and maximum values in light orange and blue. The N-terminal domain, which fitted poorly in all of the predictions (as indicated by the highlighted region), needed significant movement during refinement and is shown in **(C)** for model 1 from PEZYFoldings (group 278). Plots and 3D structures are in orange for refined models, in gray for reference structures and in blue for predictions.

**Fig. 5:**
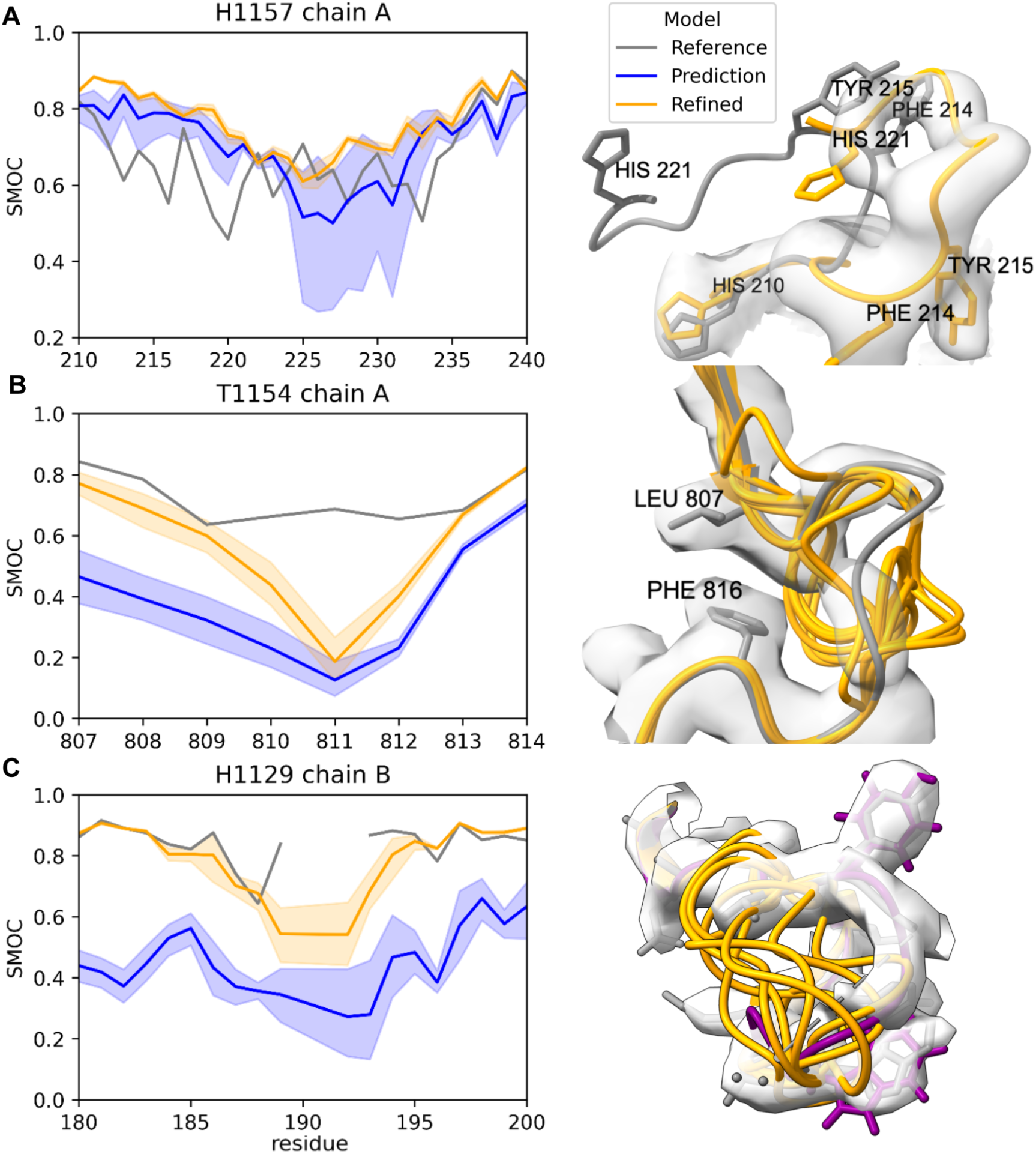
Protein loop case-studies. In all the visualizations, the target model is gray and the predictions are blue and orange before and after refinement, respectively. The dark orange line in the plot is the mean SMOC score, with the shaded region representing the minimum and maximum value for the set of predictions. **(A)** The reference model (for H1157) had a large poorly modeled loop in chain A as indicated by the low SMOC scores in 210-230 region. The best-refined predictions were a much better fit. In orange, a refined prediction from McGuffin (group 180). **(B)** This short loop, in T1154 was not modeled well enough by any predictions to be refined into the density. The low-intensity density may also be an indicator that this region is disordered. **(C)** Residues 190-191 of chain B were not modeled in the target of H1129 indicated by the dotted line. None of the predictions were able to produce a refinable loop that matched the DeepEMhancer-sharpened map in this region. However, the model submitted by Wallner (group 037), colored purple, was visually the best fitting after refinement, with side-chains at 187 and 192, well positioned.

Overall MolProbity scores, which are a log-weighted combination of the clash score, percentage of unfavoured Ramachandran dihedrals and unfavorable side-chain rotamers, generally improved after refinement with scores less than 2.0 being common. However, for a number of targets, the MolProbity scores were worse. In these cases (H1129, H1185, T1154), the provided maps had been pre-processed using DeepEMhancer ^29^ or sharpened.

Most of the best predictions based on chain ranking were improved after refinement, generally exceeding the cutoffs needed to be considered ‘accurate’ **(Table 3)**. However, some scores for T1154 and T1121o were worse after refinement due to distortions. In the case of T1154 an incorrect interaction at the N-terminus caused a poor set of rigid-bodies to be generated during refinement. In the case of T1121o, a domain was rotated perpendicular to the density causing it to fail to refine.

#### 3.2.2 Analysis of loop predictions

Given that overall the predictions were very accurate for proteins and that the top predictions required very little refinement in order to fit well into their corresponding target cryoEM maps, we decided to focus next on examining how well the loops in the top predictions were refined. Below are specific targets where the accuracy of loops was examined in detail.

#### H1157 – Complex of CtEDEM and CtPDI1P at 3.3 Å resolution

This target consists of two proteins, each with multiple domains. These were modeled in a challenging experimental map with varying resolutions. Initial inspection of the target indicated it had some modeling issues: many aromatic side-chains were not well fit to the density and a number of loops were in regions of the map that had resolution too low to be modeled with confidence. To our surprise, the best predictions modeled a loop in chain A between residues 210-230 much better than in the target and improved further upon refinement **(Fig. 5A)**. There was sufficient resolution to be confident in the modeling including density for a number of aromatic side-chains. Despite the excellent performance in modeling this large loop, predictors did a worse job at modeling a number of other loops in the target.

#### T1154 – S-layer protein A (SlaA) at 3.0 Å resolution

Many bacteria and archaea have a protein-based barrier which encapsulates the cell known as an S-layer. This target was the recently modeled outer S-layer component of the archaea *Sulfolobus acidocaldarius* ^30^. Generally, its domains were well predicted, with refined models better fitting the experimental data. Despite the overall high-resolution, a short loop between residues 810-814 had very poor density. Predictions were unable to produce loops close enough to the correct geometry to be refined into the map (**Fig. 5B)**. Although automated refinement starting from these models was not possible, the general lack of consensus amongst the predictions likely reflected some degree of disorder which was mirrored by the poor resolution seen in this region of the map.

#### H1129 – The bacteriophage pb5 protein in complex with FhuA at 3.1 Å DeepEMhancer map

Much like the swift adoption of deep-learning methods in the structure prediction community, deep-learning has been transforming image processing and reconstruction methods in the cryoEM scene. Here, a dimeric complex of the bacteriophage pb5 protein and its binding partner (the bacterial outer-membrane protein FhuA) is derived from a map which had been sharpened using the deep-learning tool DeepEMhancer ^29, 31^. Despite the overall high resolution of this map, residues 190 and 191 of a short loop were not modeled in the target structure with density dropping out in this region. Similar to the short loop in T1154, none of the predictions gave a “refineable” or even visually plausible fit **(Fig. 5C)**. However, the model provided by Wallner (group 037) was by visual inspection close and could potentially be locally fitted and refined using interactive tools such as Coot ^32^ or ISOLDE ^33^. Despite often making visual interpretation easier, an unfortunate side effect of DeepEMhancer is that lower-resolution regions of the map tend to be removed. It is possible that the unprocessed map (which we did not have) may have offered better information about this likely disordered region.

#### 3.2.3 Analysis of side-chain predictions

To examine how well CASP predictors can now predict side chains, we analyzed the side chains of predictions for target H1114 using its high-resolution 1.52 Å resolution map. The H1114 target is a hydrogenase isolated from *Mycobacterium smegmatis* that forms a large oligomeric complex formed from multiple copies of the HucS, HucL and HucM proteins ^34^. The SMOC scores for backbone and sidechain atoms of unrefined predictions compared against those of the target for each residue are shown in **Fig. 6**. Sidechain SMOC scores (SMOCs) were clearly not predicted as well as the backbone scores (SMOCb), suggesting poor atom placement **(Fig. 6A)**. An example is model 1 from Yang (group 439). In this case, although the backbone was relatively well fitted (average SMOCb=0.72), some side chains were incorrectly positioned, such as those of GLU15 and HIS166 **(Fig. 6B)**.

**Fig 6.**
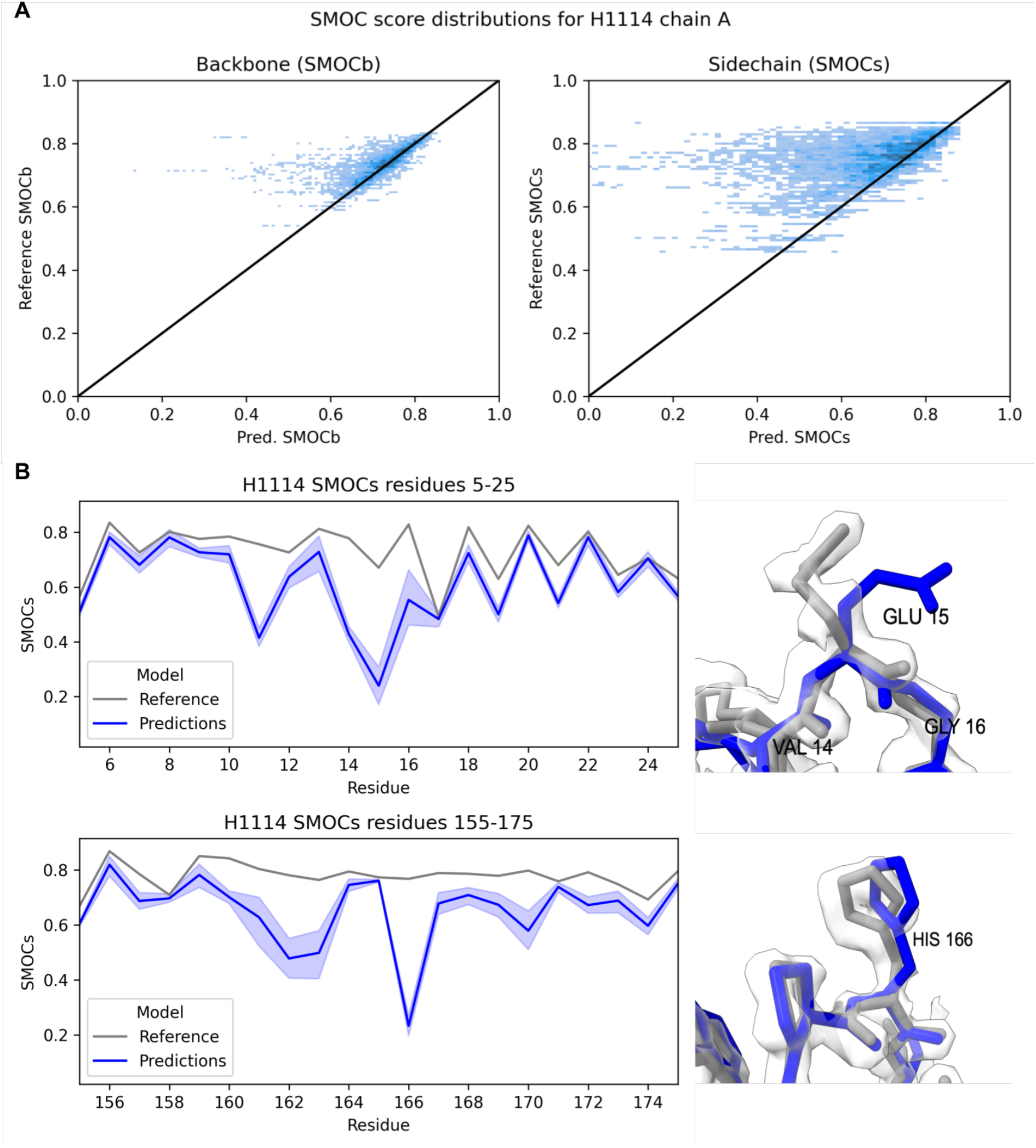
Side-chain analysis of H1114. SMOC scores for backbone and sidechain atoms of H1114 predictions compared against those of the target reference structure for each residue **(A)**. Backbone SMOCb scores (left) and sidechain SMOCs scores (right) of the reference structure vs the predictions. In **(B)** incorrectly positioned side-chains of GLU15 and HIS166 from model 1 prediction by Yang (group 439) (blue) compared to the reference (gray). These residues were consistently poorly placed by predictors.

#### 3.2.4 Refinement of T1169 – the mosquito salivary gland surface protein 1 at 3.3 Å resolution

Target T1169 is the mosquito salivary gland surface protein 1, a monomeric protein composed of more than 3000 residues involved in pathogen transmission from mosquitos. None of the predictions passed our CASP criteria for multidomain protein refinement (GDT-TS > 0.7 and LDDT > 0.7). This is potentially due to the existence of a domain in T1169 with a previously unidentified fold, and others with low sequence homology to known structures ^35^. Therefore, we decided to compare between the top-fit prediction based on chain ranking which was from Shennong (group 466), against the prediction with the highest GDT-TS score (57.7) which was from Yang-server (group 229) **(Fig. 7A)**. The Shennong model was ranked third based on GDT-TS with a score of 54.1. Note that based on global fit-to-density using ChimeraX cross-correlation (CC) scores, the Yang-server model also had a better correlation with the experimental map (CC=0.55 for Shennong and CC=0.61 for Yang-server). The refined models of each of these predictions are shown in the 3.3 Å cryoEM map **(Fig. 7A)**. SMOC scores of the predicted models show that each prediction has regions that are more accurate than the other. From the corresponding SMOC plot **(Fig. 7B)**, the CASP-criteria selected prediction produced a better refined model with a SMOC profile closer to that of the target. The poorer refinement of the Shennong group prediction **(Table 3)** is likely due to the incorrect placement of the N-terminal β-propeller towards the center of the molecule (residues 1-340), which could not be fixed during refinement **(Fig 7B)**.

**Fig. 7.**
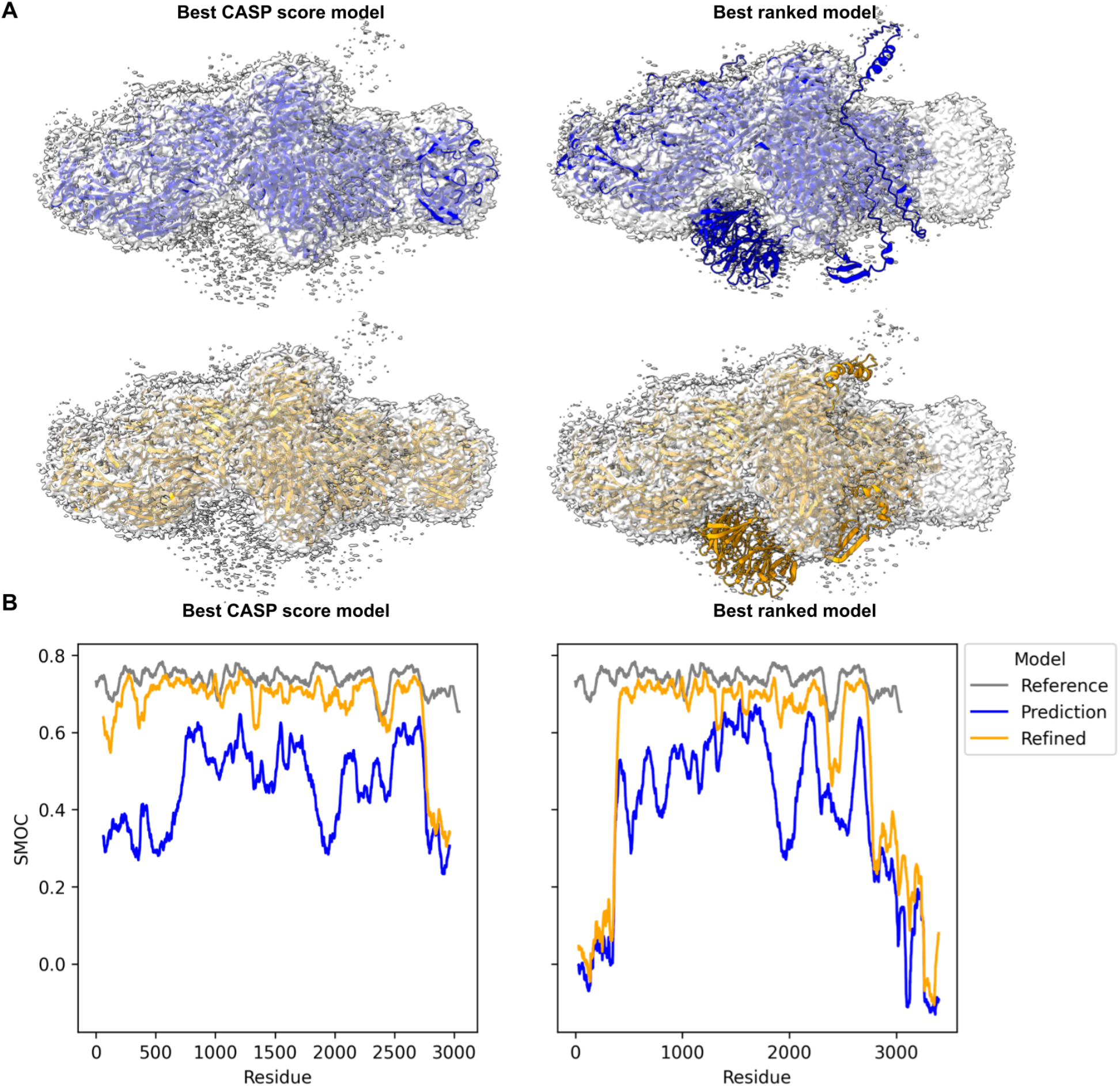
Refinement of the prediction with the best CASP score from Yang (group 229) and the best ranking Shennong (group 466). In **(A)** SMOC scores of the predictions and the corresponding structures in the experimental density. Both models have several (often different) regions which fit relatively well. The best-ranked model had more C-terminal residues modeled but had a poorly placed N-terminal β-propeller domain. After refinement **(B)**, the model based on CASP metrics had higher SMOC scores overall. The poorly placed β-propeller of the best-ranked model was too distant to be refined.

### 3.3 RNA targets: refinement of top predictions

#### 3.3.1 Selection criterion of RNA targets for refinement

Six of the eight RNA-containing cryoEM targets were selected for refinement. The two RNA-protein complexes (RT1189, RT1190) were not selected as targets due to poor prediction accuracy (RMSD>15.9Å, GDT_TS<27). A separate analysis of these predictions was performed instead ^15^. Furthermore, no predictions passed the CASP-scored selection for proteins (GDT_TS>0.7, lDDT>0.7) so we used an alternative selection process for RNA models. For each target, the previous Z_EM_ ranking was used to obtain a top 10 models which were then visually inspected to obtain a set of models we thought most likely to be refined by criteria such as limited geometric problems, and minimal chain distortions needed to move into map ^15^ **(Table 4)**. R1126, R1128, and R1149 had a single experimental structure and thus their top models by Z_EM_ were selected and after manual fitting; 6, 7, and 3 models were refined, respectively. For the three remaining RNA-only cryoEM targets, multiple experimental maps were used for refining the predicted structures.

**Table 4:** RNA predictions which were selected for refinement.

| Target | Group | Prediction model numbers |
| --- | --- | --- |
| R1126 | 232 | 1-5 |
|  | 287 | 2 |
| R1128 | 232 | 1-5 |
|  | 287 | 1,3 |
| R1136v1, R1136v2 | 232 | 1,3,5 |
|  | 287 | 4 |
|  | 325 | 1 |
| R1138v1, R1138v2 | 232 | 3,4,5 |
| R1149 | 054 | 1 |
|  | 125 | 3 |
|  | 416 | 3 |
| R1156v1 | 128 | 5 |
| R1156v2 | 128 | 5 |
| R1156v3 | 128 | 1,5 |
|  | 232 | 3 |
|  | 287 | 1 |
| R1156v4 | 232 | 3 |
|  | 439 | 2 |

For R1136, the two experimental maps, representing the ligand bound and unbound conformations, were topologically very similar, so the same models (5 total) were selected to refine into both maps. R1136 included 15 submitted models with the same RNA structure – they differed in their ligand prediction – so only 325_1 was used for refinement. For R1138, all top predictors were closest to the “mature” state, with no predictions close to the “young” state according to global topological and fit-to-map metrics. The top models (3 total) for the “mature” state were thus refined to both maps. For R1156 each map was considered separately resulting in 8 total refinements.

#### 3.3.2 Overall RNA model analysis

The RNA predictions had average SMOC scores above 0.8 after refinement for all but the young conformation of R1138 discussed below, despite predicted models starting far from the reference structure (all GDT_TS<0.7) **(Fig. 8A)**. In fact, for R1128, R1138v2, R1149, and R1156v3 targets, refined predictions surpassed the SMOC values of models fitted into the same RNA cryoEM maps as reference models **(Fig. 8A)**. Further, while prediction started with a spread of SMOC scores, the variance in SMOC score was reduced upon refinement. These results indicate that the refinement procedure was successful in fitting the models into the maps, moving all predictions to a similar solution, even in cases where large changes were needed. Compared to protein models where the fit of loops and side-chains could be assessed due to the higher resolution of the experimental maps, here the focus was on the overall fit of high level features.

**Fig. 8:**
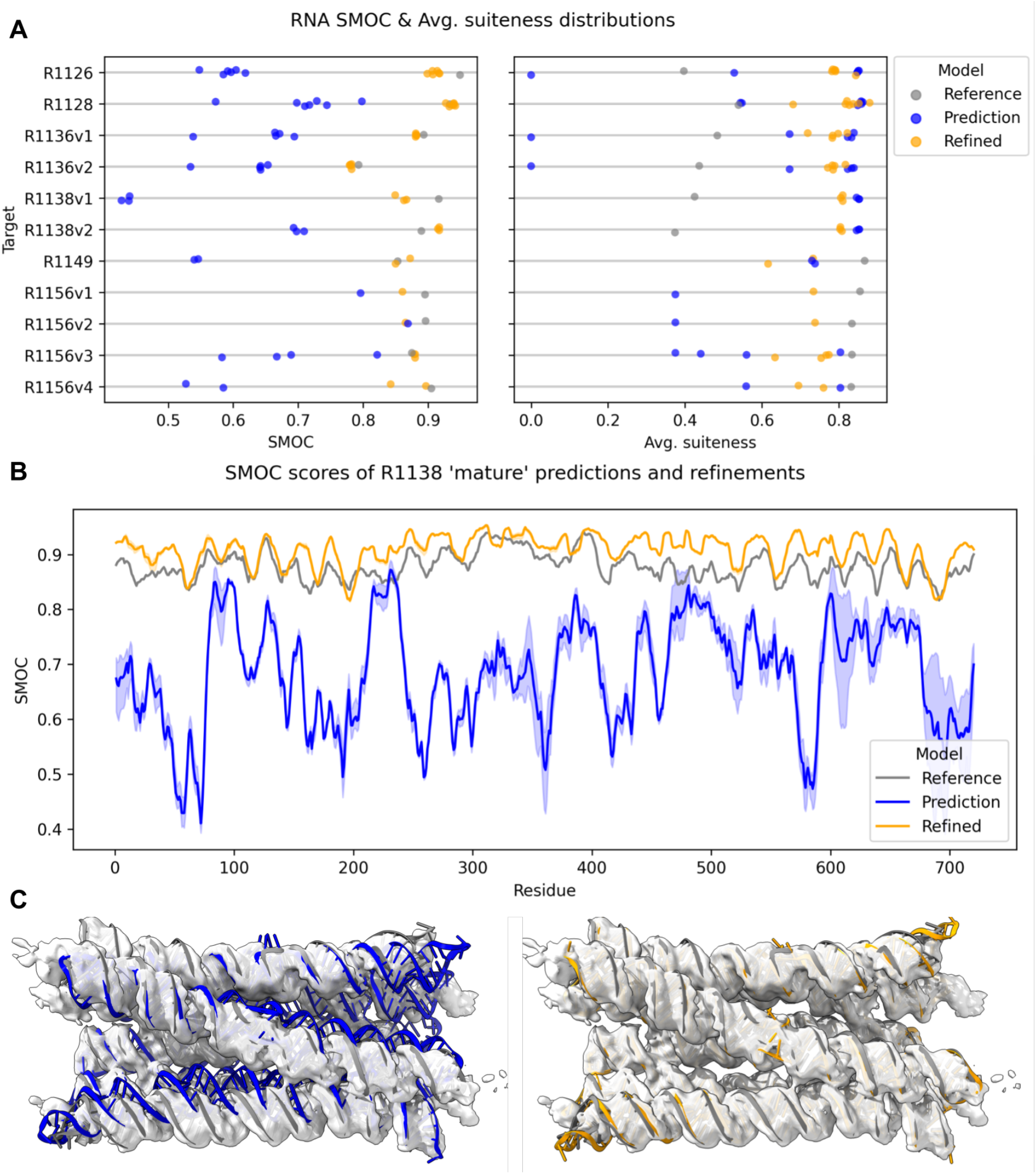
Overview of RNA refinement results. (A) The average SMOC scores for the target, predictions and refined predictions are shown alongside the RNA Validate ‘average suiteness’. **(B)** the residue level SMOC plots of R1138 in the mature conformation map and the predicted and refined models. The dark blue and orange lines are the average SMOC score for the predictions and refinements respectively, with the lightly shaded area representing the minimum and maximum values. **(C)** An R1138 prediction by Alchemy_RNA2 (group 232) in the ‘mature’ conformation map. Depicted are the prediction (blue) and refined prediction (orange) with respect to the reference model (gray).

#### R1138 a 6-helix bundle at 4.9Å resolution

A particularly interesting example for cryoEM refinement of RNA models was the predictions and refinement for R1138, a designed 6-helix bundle of RNA with a clasp (6HBC) ^36^. This target had reference structures and experimental maps for two alternative conformations, a short-lived “young” conformation and a stable “mature” conformation. The refinements for the mature conformation gave a better fit to the experimental density than the target reference structure **(Fig. 8B)** with the majority of residues having higher SMOC scores than those in the target reference structure. These predictions required significant conformational change as seen in **Fig. 8C** and **Supplemental Video 2**. The overall geometry, as assessed by the ‘average suiteness’ score (see Methods), was also better in the refined models than the reference structure **(Fig. 8A)**. However, CASP predictions for the ‘young’ conformation failed to refine to the same extent **(Fig 8A, Supplemental Video 3)**. This poorer result might be attributed to the greater degree of rearrangement of the helices and the breaking and reforming of hydrogen bonds in the kissing loop clasp required to convert from models resembling the mature conformation to the early conformation. The breaking and forming of such hydrogen bonds can in principle occur, but is unlikely, in our refinement protocol.

#### R1126 a designed “Traptamer” at 5.6Å resolution

The refined predictions of the designed RNA target R1126, a designed RNA origami scaffold for a Broccoli and Pepper aptamer pair ^37^, had lower average SMOC scores than the reference target structure. However, this result may be due to the reference structure being overfitted to the cryoEM map at the expense of realistic RNA geometry, as reflected by the low suiteness scores of the target structure compared to the refined models **(Fig. 8A)**. Selected predictions for this target had a large degree of conformational diversity with models varying between 9 and 13Å RMSD from the target. Despite our refinement protocol improving the overall fit-to-map and improving the geometry of some of these predictions, a number of predictions from Alchemy_RNA2 (group 232) exhibited an incorrect crossover between strands **(Fig. 9A)**. Fixing such issues would require breaking and rebuilding chains which is not allowed in our refinement protocol.

**Fig. 9.**
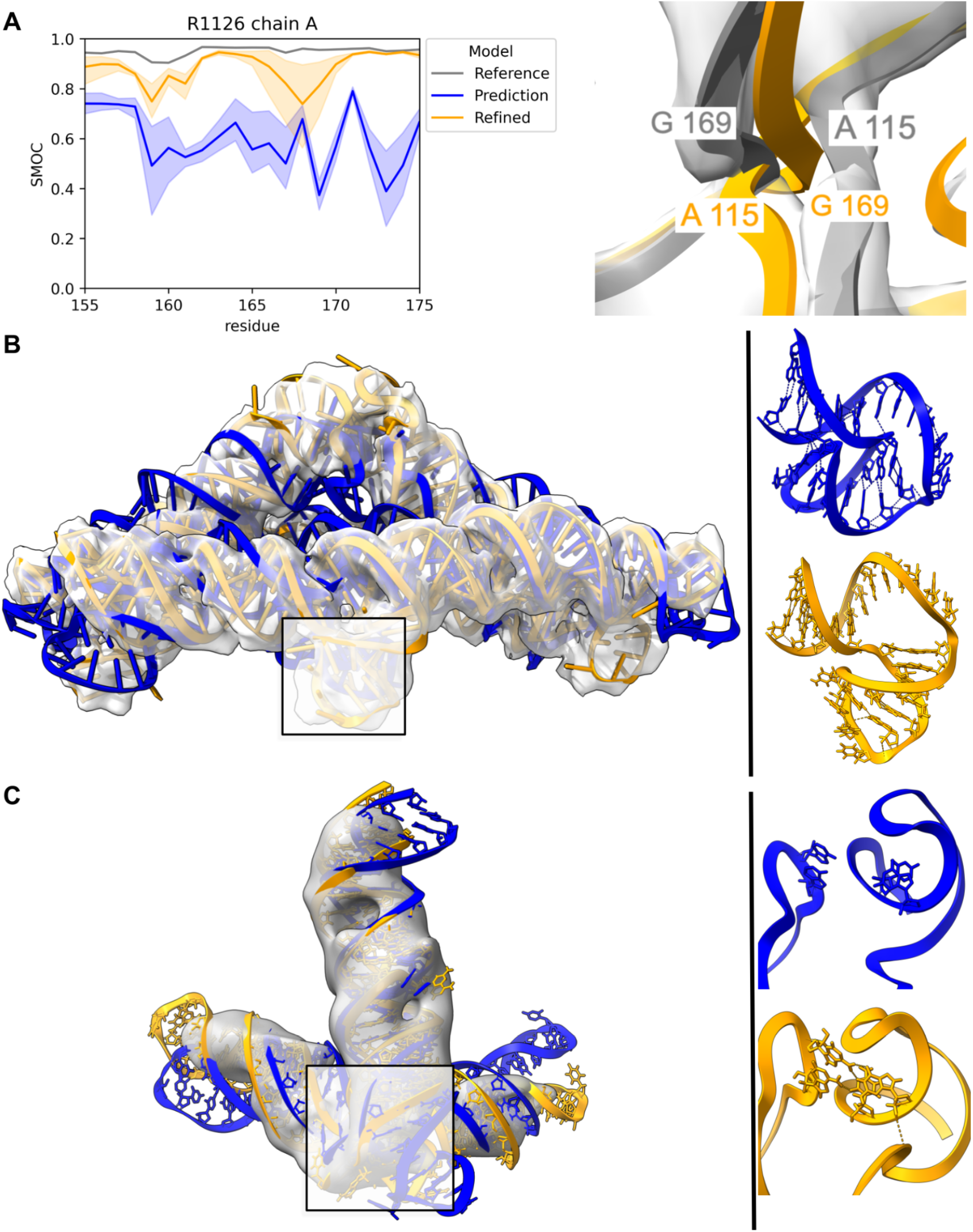
RNA refinement case studies. In all the visualizations, the target model is gray and the predictions are blue and orange before and after refinement, respectively. The dark orange line in the plot is the mean SMOC score, with the shaded region representing the minimum and maximum value for the set of predictions. **(A)** A SMOC plot of R1126 predictions and their refinements. Some refinements had residues between 155-175 with a variable SMOC score, large shaded region. This was due to strands crossing over, in some of the predictions, as shown in the right panel. **(B)** A model of R1126 from Chen (group 287) and its refinement. Overall, the R1126 predictions were refinable despite large conformational changes often being required. On the right, a close-up of the highlighted area showing the breaking of loop interaction during refinement. **(C)** A model of R1156 from Alchemy_RNA2 (group 232) and its refinement. After refining the model into the third conformation map, it better fitted the experimental density. On the right, a close-up of the highlighted area showing the formation of new interactions between an apical loop and an internal loop.

Both Alchemy_RNA2 (group 232) and Chen (group 287) provided a number of predictions which offered excellent refined models. All of these predictions required significant conformational changes to fit the experimental map. Often these movements involved breaking predicted interactions. One striking example is in the second prediction from Chen (**Fig. 9B, Supplemental Video 1)**. In the prediction, a stem-loop was curled around and interacting with an upstream helix. In order to fit the density, the stem-loop interaction was broken allowing it to move into density.

#### R1156v3 – BtCoV-HKU5 SL5 at 7.6Å

Maps and reference structures for four alternative conformations of the SL5 domain from 5’UTR from the Bat coronavirus BtCoV-HKU5 were provided for assessment in this CASP. This domain is known to have a conserved secondary structure in many coronaviruses ^38, 39^ which is thought to be important in the packaging of viral particles during infection ^40^. Maps for this target varied in resolution from 5.6 to 7.6 Å. The four refined predictions for the third conformation (R1156v3) exhibited average SMOC values slightly higher than the reference structure. Although the suiteness scores for the refined predictions were lower than for the reference structure, in all but one case they were better than the unrefined predictions. In contrast to the Traptamer example above, where refinement involved the breaking of an interaction of a apical loop, the refinement of the second prediction from Alchemy_RNA2 involved the formation of an interaction between a apical loop and an internal loop in another part of the model **(Fig. 9C)**.

## Discussion

In CASP15, 29% of the total targets, 67% of the RNA-containing targets, were determined using cryoEM. The accuracy of predictions for protein targets assessed in this paper and the overall quality of experimental maps allowed many predictions to be further refined to near-native conformations. Compared to most CASP assessments, where a single reference model has been used as the ground truth, cryoEM assessment finds itself in a privileged position. To aid the assessment, CryoEM maps are typically available in conjunction with target reference models – which are after all just best attempts at model building using the experimental map, human knowledge and current state of the art technology. This is particularly important, as cryoEM data tends to have lower resolutions than crystallographic experiments. Because 3D reconstructions are built from averages of many particles, they may also capture continuous motions and flexibility of the visualized macromolecule, which can then manifest itself as lower resolution regions. There is thus an added degree of uncertainty in any static 3D structure that is derived from cryoEM data.

One model, which particularly highlighted the importance of experimental data this year, was H1157. This model had an average resolution of 3.3 Å with many regions of the map having lower local resolution. Intriguingly, a large loop which was erroneously modeled in the target was much better modeled by the best predictions, fitting the density with aromatic side chains well placed. If, on the other hand, we only had the target model as ground truth (i.e., if we did not use the experimental map for assessment), these better predictions would have not been observed.

For the majority of targets, where the author’s submitted model (target reference model) and experimental map were in good agreement, some parts of the predicted models resulted in better fit to map following refinement. At the same time, many targets had loops, which were not predicted so well, often surprisingly short. Typically, the geometry of these loops varied amongst predictions, with many failing to be refined because they were too distant from the target. The lack of consensus amongst some of these loops was often reflected by lower local resolutions in the experimental map. While we did not investigate the relationship between these two phenomena in this paper, in CASP14 cryoEM assessment, we showed anticorrelation between the standard deviation of the SMOC scores of the predicted models (SMOC SD) and SMOC scores of the target structures ^2^.

The strong correlation between the rankings based on the cryoEM-based docking score and the composite S_CASP15_ score shows that high quality models can often be picked using experimental data alone. For model building practitioners, this is particularly relevant, as reference structures may not be available. Given the difficulty of building models into experimental maps and the fact that there isn’t a single prediction tool which excels across all targets, docking and ranking offers an approach to screen for good starting models, potentially from multiple structural prediction tools.

Unfortunately, some maps provided by the experimentalists had already been sharpened with DeepEMhancer ^29^. This caused a degradation in MolProbity scores, likely because the TEMPy-REFF GMM puts more weight on the sharpened map, overpowering the geometry restraints. Another unfortunate side-effect of DeepEMhancer maps is that low-resolution regions tended to disappear entirely in the sharpened maps. Many of the predictions displayed a diverse set of loops in these regions. While sharpened maps may aid in model building, low-resolution regions can be an important indicator of flexibility and disorder. In future CASP cryoEM assessments it would be useful to encourage the authors to provide unsharpened maps, and even half maps for further assessments.

For the first time in CASP history, RNA structures were provided as targets and the majority of them had associated cryoEM density maps. Compared with the proteins, these RNA maps had much lower resolutions. Indeed, in some maps such as those of R1156, pitches of helices were not always visible. Local fit-to-map scores, such as the newly developed SMOCr, can aid the assessment of RNA models in these challenging resolutions. Here, this local fit analysis indicated that many secondary structures and important geometric features can be accurately predicted. Furthermore, we showed that *in silico* models can, after further refinement, offer plausible models that better reflect the experimental maps even at low resolutions. However, at such low resolutions, it is possible for many alternative structures to fit the density with equal likelihood. Due to both the known flexibility of the RNA molecules and the heterogeneity of the experimental maps, ensembles of models are arguably a more accurate way to describe the underlying experimental data ^15, 41^.

Despite the overall quality of predictions, some reorientation of domains and secondary structure elements was often required, particularly for RNA models. The multistage pipeline presented offers an approach to fitting and refinement of structural models into cryoEM maps at a variety of resolutions. The use of progressively smaller rigid-bodies has been shown to aid the fitting of models that require large conformational changes ^16^. However, if the models contain topological errors or significant misplacements of elements even such a detailed approach will fail.

As mentioned above, in CASP15 there were two RNA-protein complexes (RT1189, RT1190). The predictions associated with these targets were not refined due to poor accuracy ^15^. Given the current progress in the structure prediction field, we expect further improvement on this front in future CASPs.

CryoEM has been an important method for elucidating large atomic structures, albeit often at a lower resolution than crystallographic experiments. This CASP15 for example, the largest monomeric structure in the history of CASPs, T1169, was a cryoEM target. Moreover, cryoEM experiments are now not just capturing large molecules but often achieving atomic levels of detail. In CASP15, focussed maps for the target H1114 reached an astonishing resolution of 1.52 Å. While at such resolutions, computational models are not required for model building, high-resolution data offers an opportunity to assess accuracy at an even finer level. Using the SMOC score separately for backbone (SMOCb) and side-chains (SMOCs), allowed us to show that while the overall backbone geometry of H1114 predictions was well modeled, sidechain orientations did not always agree with the experimental map. Given the progress in both protein structure prediction and cryoEM fields, we foresee such analyses becoming more routine in the future.

## Supporting information

Refinement of R1126TS287_2

Refinement of R1138TS232_3 in mature conformation map

Refinement of R1138TS232_4 in young conformation map

## Acknowledgements

This research was supported by the cooperation of Leibniz Institute of Virology (as part of Leibniz ScienceCampus InterACt, funded by the BWFGB Hamburg and the Leibniz Association) and by the Strategic Incentive Program of LIV and the Landesforschungsförderung Hamburg (Hamburg-X) (to T.M, J.G.B, and M.T). The PhD studentship of L.E. is co-funded by the Collaborative Computational Project for Electron cryo-Microscopy (CCP-EM). This research was supported by the US National Institute of General Medical Sciences (NIGMS, National Institutes of Health) grants R01 GM100482 (A.K.) and R35 GM122579 (R.D.); Stanford Bio-X (to R.D. and R.C.K.); and the Howard Hughes Medical Institute (HHMI, to R.D.). This article is subject to HHMI’s Open Access to Publications policy. HHMI lab heads have previously granted a nonexclusive CC BY 4.0 license to the public and a sublicensable license to HHMI in their research articles. Pursuant to those licenses, the author-accepted manuscript of this article can be made freely available under a CC BY 4.0 license immediately upon publication.

## Author contributions

T.M. and M.T coordinated the research. T.M. performed the cryoEM fitting and refinement of the protein and RNA predictions, and analyzed the results together with M.T.. L.E. and D.R. designed, performed and analyzed the protein docking and ranking experiments. R.C.K. and R.D. performed analyses of RNA predictions and contributed to the design of the RNA refinement protocol. T.M., J.G.B. and M.T. developed the protein and RNA refinement protocol. A.K. and J.G.B. contributed to the analysis of protein targets. T.M. and M.T drafted the manuscript with contributions from all authors.

## Listings

**Listing 1:**
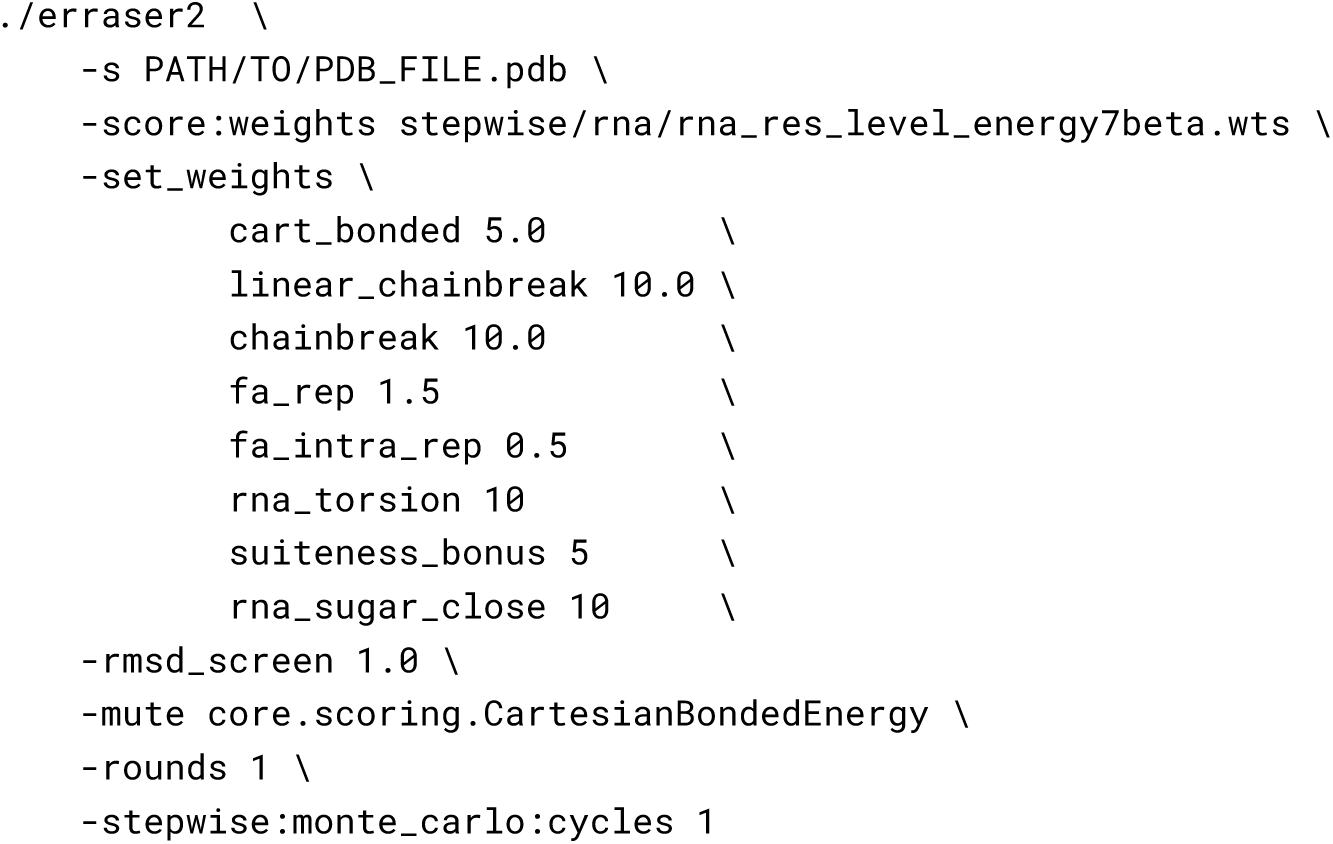
ERRASER2 command. The following command was used to run ERRASER2 using the nightly version (2021.16.61629) of Rosetta.

## Supplementary

**Supplemental Video 1:** Refinement of R1126TS287_2 **Supplemental Video 2:** Refinement of R1138TS232_3 in Mature map **Supplemental Video 3:** Refinement of R1138TS232_4 in Young map

